# Idiosyncratic genome degradation in a bacterial endosymbiont of periodical cicadas

**DOI:** 10.1101/199760

**Authors:** Matthew A. Campbell, Piotr Łukasik, Chris Simon, John P. McCutcheon

## Abstract

When a free-living bacterium transitions to a host-beneficial endosymbiotic lifestyle, it almost invariably loses a large fraction of its genome [1, 2]. The resulting small genomes often become unusually stable in size, structure, and coding capacity [3-5]. *Candidatus* Hodgkinia cicadicola (*Hodgkinia*), a bacterial endosymbiont of cicadas, sometimes exemplifies this genomic stability. The *Hodgkinia* genome has remained completely co-linear in some cicadas diverged by tens of millions of years [6, 7]. But in the long-lived periodical cicada *Magicicada tredecim*, the *Hodgkinia* genome has split into dozens of tiny, gene-sparse genomic circles that sometimes reside in distinct *Hodgkinia* cells [8]. Previous data suggested that other *Magicicada* species harbor similarly complex *Hodgkinia* populations, but the timing, number of origins, and outcomes of the splitting process were unknown. Here, by sequencing *Hodgkinia* metagenomes from the remaining six *Magicicada* species and two sister species, we show that all *Magicicada* species harbor *Hodgkinia* populations of at least twenty genomic circles each. We find little synteny among the 256 *Hodgkinia* circles analyzed except between the most closely related species. Individual gene phylogenies show that *Hodgkinia* first split in the common ancestor of *Magicicada* and its closest known relatives, but that most splitting has occurred within *Magicicada* and has given rise to highly variable *Hodgkinia* gene dosages between cicada species. These data show that *Hodgkinia* genome degradation has proceeded down different paths in different *Magicicada* species, and support a model of genomic degradation that is stochastic in outcome and likely nonadaptive for the host. These patterns mirror the genomic instability seen in some mitochondria.

## Results

Like all sap-feeding insects, cicadas depend on specialized endosymbiotic microorganisms for supplementation of their nutrient-poor diet of plant sap [9-11]. One of these microbes, an alphaproteobacterium called *Hodgkinia*, is associated with all reported cicadas [10]. As is typical for bacterial endosymbiont genomes, *Hodgkinia’s* genome is extremely reduced (∼150kb at its largest in some cicadas), rendering it completely dependent on its cicada host and partner bacterial endosymbiont *Sulcia* for basic biological function [4]. Despite a very high rate of sequence evolution [8], *Hodgkinia* genomes can be remarkably stable in structure and gene content, changing little in cicadas diverged by tens of millions of years [7, 12]. In other cicadas, *Hodgkinia* has evolved into two or more distinct but codependent genomic and cellular lineages that are present in individual hosts, which have undergone reciprocal gene inactivation and loss [7]. We refer to this unusual process as “splitting”. We have shown that in the long-lived periodical cicada *Magicicada tredecim, Hodgkinia* has split into dozens of small distinct genomic circles encoding few recognizable genes [8]. Here we analyze the *Hodgkinia* metagenomes of the six remaining *Magicicada* species in order to understand the timing and outcome of this splitting process across the cicada genus.

### *Hodgkinia* is complex in all *Magicicada* species

Our new sequencing data confirm [8] that *Hodgkinia* is comprised of many distinct genomic circles in all species of *Magicicada* (Fig. 1, Tables 1, S1). We refer to individual circular-mapping *Hodgkinia* genomic contigs as ‘circles’ because, though we know some reside in distinct cells [8], we currently do not know whether most of these molecules are chromosomes that share the same cell or genomes representing different cell types [8]. We refer to the total complement of *Hodgkinia* contigs assembled in a single species of cicada—whether closed into circles or not—as that species’ *Hodgkinia* Genome Complex (HGC). The smallest HGC is found in *M. tredecula* and consists of at least 153 contigs totaling 1.20 megabases (Mb) of DNA, while the largest is from *M. neotredecim* and consists of 215 contigs totaling 1.58 Mb (Table 1). In each *Magicicada* species, we identified between 26 and 42 contigs with large-insert mate pair data suggesting they were circular DNA molecules. We were able to fully close these contigs into circular molecules in at least 20 instances in all cicada species (Fig.1, Table 1). Contigs with mate pair, PCR, and/or Sanger sequence data supporting their circularity were considered putative circles if they were not fully closed (Fig. 1). The combination of confirmed and putative circular molecules comprised between 51.4% and 72.5% of the total DNA in each HGC (the remaining contigs lack end-joining data; Fig. 1). Individual completed circles range in size from 0.69kb to 70.5kb, contain a maximum of 27 genes, a minimum of 1 gene (with a single exception, a circle encoding only a single pseudogene), and span as much as a 653-fold range of sequencing coverage (Table S1). There is an even higher range in coverages for contigs that did not assemble into circular molecules: we find that contigs from *Magicicada* HGCs span at least a 2,500-fold difference in average sequencing coverage in each *Magicicada* species (Table 1).

**Figure 1:**
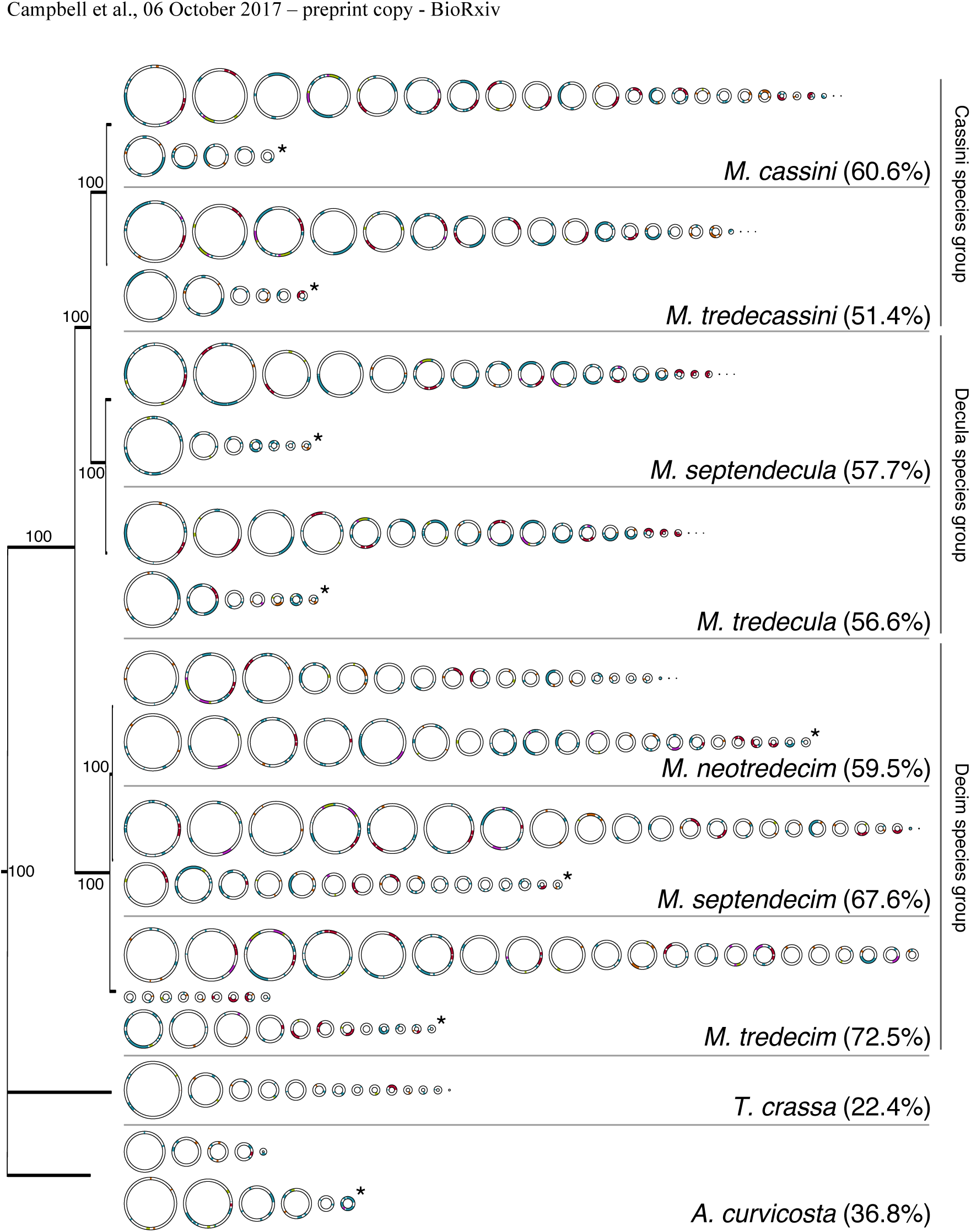
*Hodgkinia* genomic complexity in all study species. Left: Phylogeny of the cicada species used in this study, based on all 13 protein-coding and both ribosomal genes from the mitochondrial genomes. The cicada *Diceroprocta semicincta* was used to root the tree, but was not included in the figure. Bootstrap support values are shown on each resolved node. Right: Diagrams representing the confirmed and putatively circular molecules of the HGC in all study species. Rows with an asterisk at the end represent putative circular molecules. On each circle, red regions represent rRNA genes, green represents histidine synthesis genes, orange represents cobalamin synthesis genes, purple represents methionine synthesis genes, blue represents all other genes, and white space represents noncoding DNA. Values in parentheses indicate the proportion of total *Hodgkinia* DNA from each cicada species represented by the circular molecules. The three species groups are annotated next to the species labels.

**Table 1.** Summary statistics for all HGCs described in this work. Total HGC size is a sum of all *Hodgkinia* contigs no matter whether they are closed into circular molecules or not. The number of unique genes found in other *Hodgkinia* genomes range from 168-183.

| Species | Species abbreviation | Number of contigs | Total HGC size (Mb) | Total # of circles | Cumulative size of circles (Mb) | Unique genes | Total genes | Fold-coverage difference |
| --- | --- | --- | --- | --- | --- | --- | --- | --- |
| *M. cassini* | MAGCAS | 118 | 1.27 | 29 | 0.77 | 142 | 306 | 6,376 |
| *M. tredecassini* | MAGTCS | 201 | 1.42 | 26 | 0.73 | 145 | 316 | 5,494 |
| *M. septendecula* | MAGSDC | 119 | 1.22 | 27 | 0.71 | 139 | 297 | 4,827 |
| *M. tredecula* | MAGTDC | 153 | 1.20 | 27 | 0.68 | 140 | 318 | 3,189 |
| *M. neotredecim* | MAGNEO | 215 | 1.68 | 41 | 1.00 | 139 | 333 | 3,379 |
| *M. septendecim* | MAGSEP | 166 | 1.64 | 39 | 1.11 | 137 | 314 | 5,723 |
| *M. tredecim* | MAGTRE | 118 | 1.53 | 42 | 1.11 | 135 | 305 | 2,500 |
| *T. crassa* | TRYCRA | 106 | 1.16 | 14 | 0.26 | 137 | 203 | 947 |
| *A. curvicauda* | ALECUR | 138 | 0.95 | 11 | 0.35 | 136 | 199 | 830 |

We identified between 135 and 145 of the 186 unique protein- and RNA-coding genes annotated as functional in at least one of *D. semicincta* and *T. ulnaria* in each *Magicicada* HGC [7, 12]. In all species, several additional genes were identified as truncated fragments or obvious pseudogenes. Because of the very low coverage of some contigs and the extremely rapid rate of *Hodgkinia* sequence evolution [8], it it is likely that many of the remaining genes are present but were either not fully assembled or are not recognizable due to their low sequence similarity to other annotated *Hodgkinia* genes.

We also sequenced *Hodgkinia* from two cicada species that are closely related to *Magicicada* [13, 14] (Fig. 1, Table 1). The HGCs from Australian cicada species *Aleeta curvicosta* and *Tryella crassa* are similar in many ways to HGCs from *Magicicada*, but somewhat less complex. However, we generated less sequencing data for these species (∼8.7 gigabases (Gb) for *A. curvicosta*, ∼1.5 Gb for *T. crassa*, compared with an average of ∼30.5 Gb per species of *Magicicada*) and so this relative simplicity may be due to sequencing effort. We therefore primarily focus our analyses on the *Hodgkinia* genomes from *Magicicada* species. Nevertheless, data from these outgroup species allowed us to infer whether *Hodgkinia* lineage splitting began in *Magicicada* or whether independent *Hodgkinia* lineages existed before *Magicicada* diverged from its common ancestor.

### The origin of *Hodgkinia* lineage splitting predates the diversification of the genus *Magicicada*

To determine whether *Hodgkinia* lineage splitting started in the *Magicicada* genus or predated its origin, we reconstructed phylogenetic trees for *Hodgkinia* genes present in multiple copies in at least one of the two outgroup species. If splitting began independently in each genus, we would expect phylogenetic trees inferred from individual *Hodgkinia* genes to be monophyletic within each cicada genus (Fig. S1A). However, if splitting predated the divergence of *Magicicada* from *Tryella/Aleeta*, then gene trees should show two or more strongly supported monophyletic clades, each consisting of copies of genes from *Magicicada* along with *Tryella* and *Aleeta* (Figs. S1B-D). Though we see some cases where gene trees form monophyletic groupings within cicada genera (Fig. S1A), we also find several instances where gene phylogenies reveal two (Fig. S1B-C) or three (Fig. S1D) well-supported clades that group *Magicicada* genes with at least one gene copy from *Tryella* or *Aleeta*. It is possible to see both patterns because not all redundant genes from split lineages are retained in the new lineages [7]. Overall these patterns show that lineage splitting in *Hodgkinia* began before the *Magicicada, Tryella,* and *Aleeta* diverged from one another. We estimate that the last common ancestor of these genera had a minimum of three *Hodgkinia* lineages (Fig. S2A-C), similar to the complexity of *Hodgkinia* in the cicada *Tettigades undata* [7].

### *Hodgkinia* lineage splitting is ongoing in *Magicicada* species groups

Having found evidence that *Hodgkinia* splitting had started prior to the divergence of *Magicicada* from its ancestor with *Tryella* and *Aleeta*, we tried to assess whether most circular molecules were formed prior to the diversification of *Magicicada* and were conserved throughout the genus, or if lineage splitting is a process that has been ongoing since the origin of *Magicicada*. If most lineage splitting occurred in the ancestor to all *Magicicada*, then most gene copies should have representatives in all extant cicada species. If most lineage splitting occurred within species groups, then many gene copies should be unique to species groups and not shared throughout the genus. Gene phylogenies generated with six representative *Hodgkinia* genes show multiple but relatively few well-supported clades with representatives of all three *Magicicada* species groups (Fig. S2). In some cases, we identified only one gene copy per HGC, with gene phylogenies that recapitulate host phylogeny (Fig. S2A). This pattern suggests that there was a single copy of the gene in the last common ancestor of *Magicicada*, and that the genomic circle it was on may not have undergone splits. In other cases, the single ancestral gene copy co-diversified with hosts, but also underwent splits in some species groups (Fig. S2B). Trees of other genes form between two and five highly supported clades that include copies from all species groups (Fig. S2C-F), showing evidence for variable amount of additional splitting after species groups diverged (Fig. S2G). These phylogenies show that a minimum of five distinct *Hodgkinia* lineages existed in the last common ancestor of *Magicicada*.

These data suggest that most of the splitting we see in Fig. 1 happened after *Magicicada* started to diversify. If this is true, we expect that the similarity of HGCs should diminish as a function of cicada phylogenetic distance. In comparing extant circular molecules between cicada species, we find few clearly homologous circles with identical gene sets conserved in all *Magicicada* species. Because comparative genomic methods are generally based on sequence similarity and synteny comparisons, and we found little obvious synteny to compare, we developed a metric based on the Jaccard Index [15] to quantify the similarity in gene content of the finished circles between cicada species. We call this metric the *Hodgkinia* Similarity Index (HSI, Fig. 2). We calculate the HSI as follows, for hypothetical circular molecules A and B (**Equation1**):

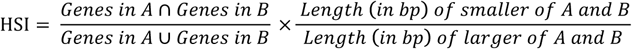

**Figure 2:**
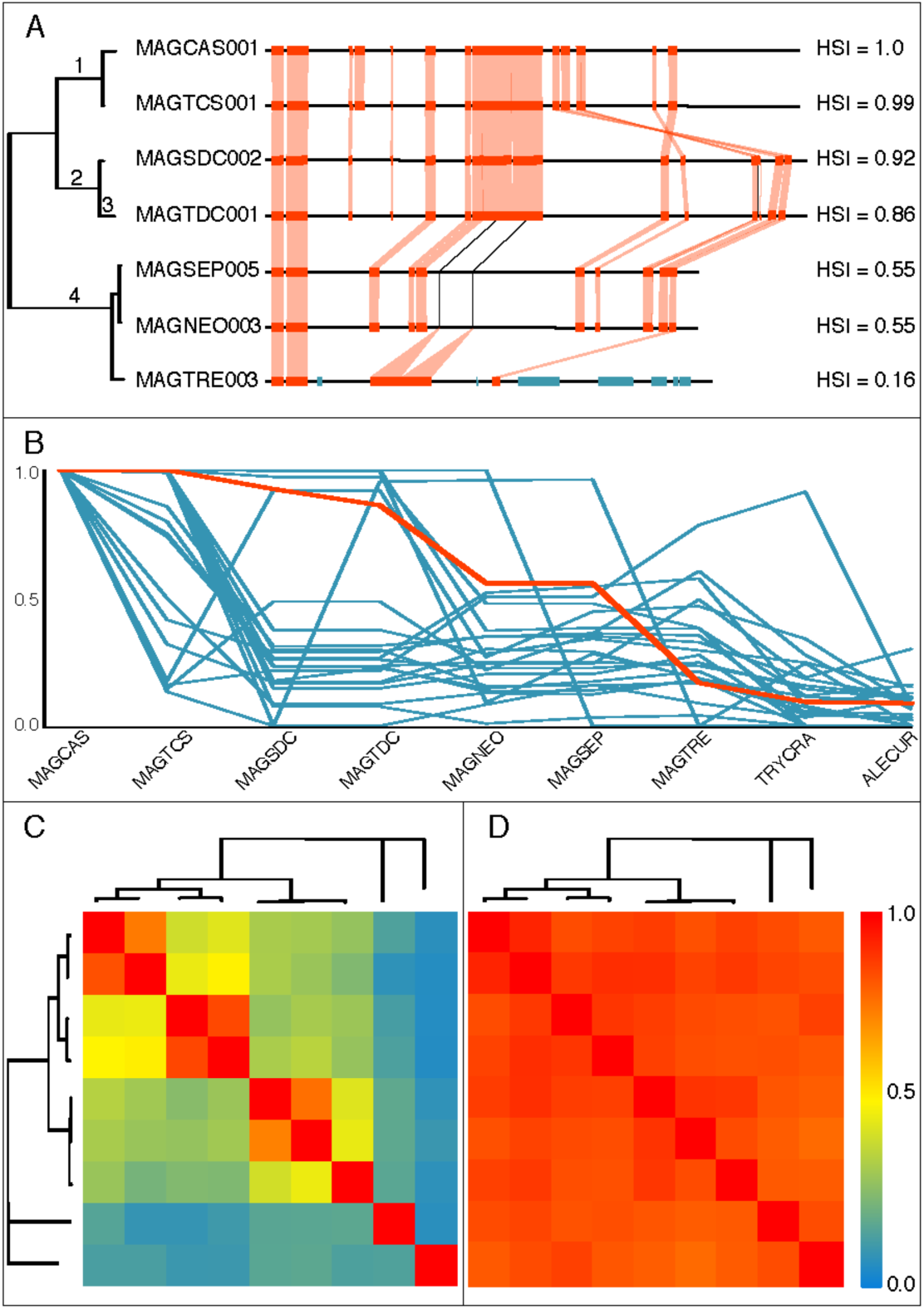
HSI scores for individual circles and HGCs. **(A)** Illustration of synteny conservation between homologous circular molecules. Shown is the reference circle MAGCAS001 and the circle most similar to it from all other *Magicicada* species (abbreviations are taken from Table 1). Horizontal black lines represent the genome backbone, orange boxes are genes shared between a given circular molecule and MAGCAS001. Blue bars represent genes present in a given circular molecule but not present on MAGCAS001. Shaded vertical lines show gene homologs present on different circles, black lines connect putative homologs over gaps in some genomes. HSI scores between the reference and all circular molecules are shown on the right. Numbers on the phylogenies represent inferred mutational events on the respective lineage: (1) genome rearrangement, (2) and (3) individual gene loss events, (4) loss of five genes. The exact branch on which (4) occurred is ambiguous. Three contigs from *M. tredecim* seem to be homologous to the reference circle when joined together, but we could not close them to a single circle so they were not included in the HSI analysis. **(B)** Distribution of all HSI scores for *M. cassini*. The x-axis shows each species *M. cassini* was compared with, and the y-axis shows the HSI score. The bold orange line represents the circles shown in (A). **(C)** Heatmap showing pairwise average HSI scores between all species. **(D)** Heatmap showing pairwise average Jaccard Index of the whole *Hodgkinia* gene set in each species. In both B and C, a score of one indicates that the two species are identical, and zero indicates that they share no genes in common. All trees are taken from Figure 1.

Briefly, a finished circular molecule of one cicada species is compared to a circular molecule of another cicada species. We calculate the Jaccard Index of the two gene sets (the intersection of gene sets divided by the union, the left half of Eq. 1) and multiply that by the ratio of the length of the smaller circle divided by the length of the larger one (right half of Eq. 1). We calculate this pairwise value for all circles of a species and report the average HSI score between those two cicada species. We then repeat this for all pairwise comparisons of cicada species. An HSI value of one indicates the two circles have the same functional genes and are the same length, whereas a value of zero indicates they share no common genes. Because the circles have on average very low coding densities and have apparently undergone rearrangements in some cases (Fig. 2A), this metric does not take gene co-linearity into account. It is also not (necessarily) a true measure of homology since it does not distinguish between conservation of an ancestral circle and convergent evolution to a similar state. Rather it is a rough metric to score the overall similarity of HGCs between cicada species in the absence of much calculable similarity (Fig. 2B). It is also a conservative metric, since there will undoubtedly be homologous circular molecules that were not completely assembled and thus not calculated in the HSI.

We find a strong phylogenetic signal in HSI scores, where HGCs between species pairs (*M. cassini–M. tredecassini, M. septendecula–M. tredecula,* and *M. septendecim–M. neotredecim*) are highly similar to one another (0.80 HSI on average, Fig. 2C). This is expected given that each of these species pairs are estimated to have diverged from each other less than 50 thousand years each ago [13]. The HSI scores degrade quickly with increased phylogenetic distance, however. Pairwise comparisons between *M. tredecim-M. neotredecim* and *M. tredecim-M. septendecim* (500 thousand years diverged [13]), Cassini species with Decula species (2.5 million years diverged [13]), and Cassini and Decula species with Decim species (4 million years diverged [13]) give average HSI scores of 0.43, 0.46, and 0.29, respectively. This lack of similarity is remarkable given that the HSI between the single *Hodgkinia* genomes of *Diceroprocta semicincta* and *Tettigades ulnaria*, which diverged more than 60 million years ago [16-19], is 0.88.

Our combined phylogenetic and HSI analyses suggest that splitting began in the ancestor of *Magicicada, Tryella,* and *Aleeta* (into 2-3 circles), continued somewhat in the ancestor of all *Magicicada* (into at least 5-6 circles), but that splitting accelerated dramatically (into 20+ circles) after *Magicicada* began diversifying.

### *Hodgkinia’*s overall function mostly remains intact

The long-term stability of endosymbiont genomes is often attributed to the importance of their function to host survival [3, 20, 21]. Since *Hodgkinia* is clearly experiencing dramatic genomic instability, we wanted to test whether the complete ancestral *Hodgkinia* gene set was retained in HGCs in different *Magicicada* species. To directly compare gene complements between *Hodgkinia* HGCs, and to be consistent with the HSI, we calculated the Jaccard Index of each gene set for all pairwise comparisons of all *Magicicada* species. Similar to the HSI, a score of 1 would indicate that two cicada species have identical *Hodgkinia* gene sets, and a score of 0 would indicate that no genes are shared. We find that HGC gene sets within closely related species pairs are very similar (0.90 on average, Fig. 2D). Pairwise comparisons between *M. tredecim-M. neotredecim* and *M. tredecim-M. septendecim* (0.86), Cassini species with Decula species (0.87), and Cassini and Decula species with Decim species (0.86) also remain very similar, in contrast to the HSI scores calculated for these comparisons (compare Fig. 2C to 2D). These data indicate that while the patterns of *Hodgkinia* genome fragmentation is different in divergent *Magicicada* species, the overall set of retained genes is similar. For a sense of scale, the same analysis for cicadas diverged for dozens of millions of years [16-19], such as *Magicicada* and *D. semicincta, Magicicada* and *T. ulnaria*, and *D. semicincta* and *T. ulnaria* gives values of 0.82, 0.77, and 0.92, respectively. We note again that not all *Hodgkinia* genes present in *Magicicada* may have fully assembled due to the complexity of the dataset, so the true values for *Magicicada* HGCs may be higher than what we report here.

### Lineage splitting leads to different gene dosages

It seemed possible that lineage splitting in *Hodgkinia* might be beneficial for the host, perhaps as a mechanism to control the dosage of *Hodgkinia* genes. Under this hypothesis, we would expect similar gene dosages in comparisons of various *Magicicada* species. To calculate gene dosage in an HGC, we summed the average coverage of all contigs on which a given functional gene is found, scaled to the most abundant gene for each species. We find that the relative abundances of genes are similar within species groups (cicadas diverged less than 50 thousand years ago [13]), but not between species groups (Fig. 3). Principal coordinates analysis of relative gene abundances of all genes present in any species clusters the Decula and Cassini species groups together, *M. neotredecim* and *M. septendecim* together, and the remaining species – including M. tredecim separated from *M. neotredecim-septendecim* by only 0.5 Mya [13] – separately (Fig. S3A). This can be more clearly seen when only considering genes annotated in all species (Fig. S3B). This grouping is qualitatively similar to the HSI results, and suggests there is not the convergent pattern of gene dosage outcomes that might be expected if the host was dictating the process. Rather the gene dosage outcomes are stochastic and thus only similar in comparisons between very closely related cicadas.

**Figure 3:**
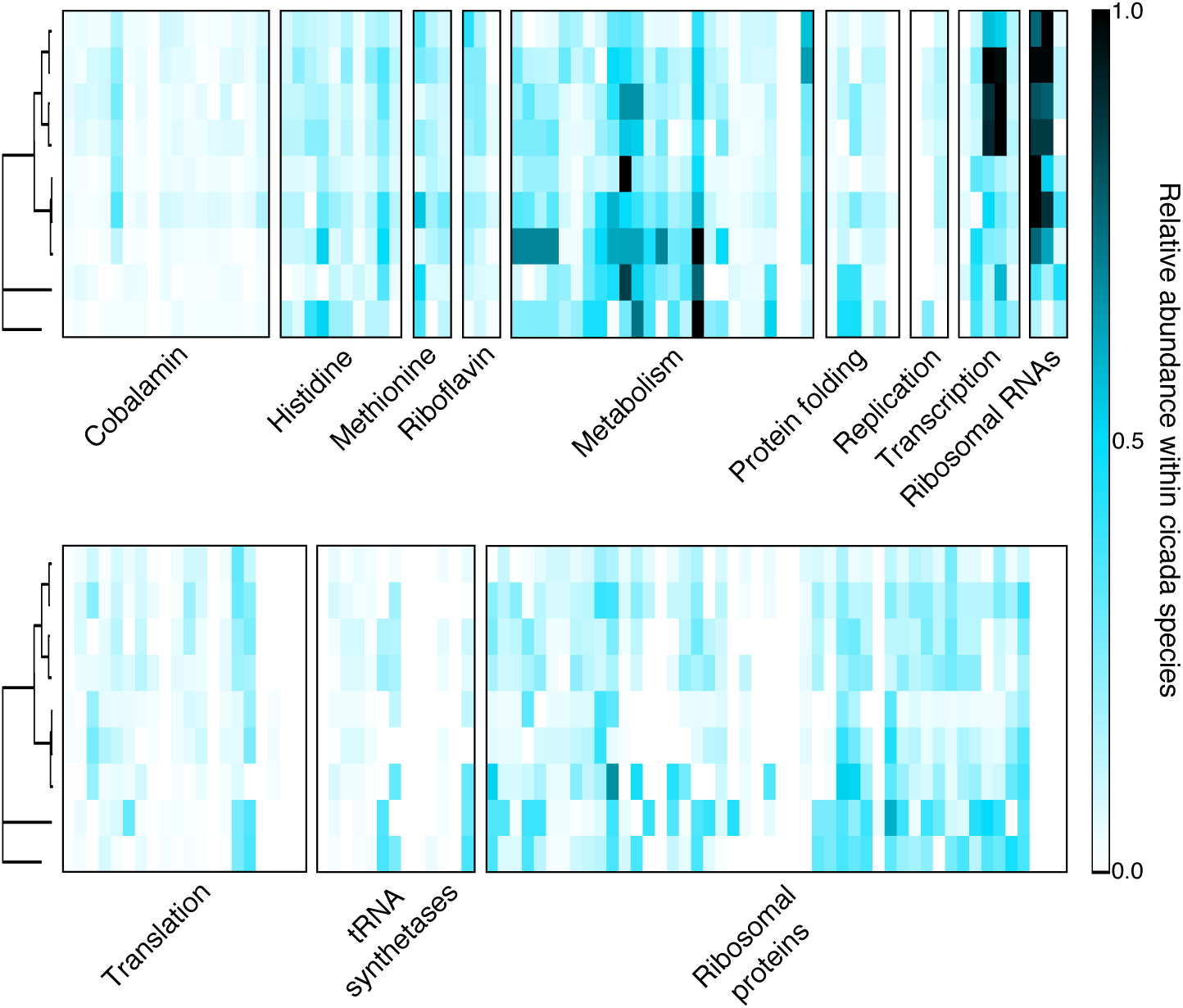
Relative gene abundance in all study species. Heatmaps showing the relative abundance of each gene in each species, ordered by gene category. A value of one (black) indicates the most abundant gene in that species, and zero (white) indicates that gene is absent in that species. Columns that are completely white represent genes that were not annotated in any species, so have either been lost, are present on broken contigs, or are present on contigs that did not otherwise assemble in our experiments. Trees are taken from Figure 1.

## Discussion

Many endosymbioses consist of two or more partners that are strictly reliant on one another for survival. The eukaryotic cell is now completely intertwined with and dependent upon its mitochondria (with one known exception [22]), and mitochondria cannot survive outside of their host cell. Similarly, cicadas require *Hodgkinia* for survival, and it is unlikely that *Hodgkinia* could survive outside of cicada cells. Despite these obligate host-symbiont co-dependencies, each partner can experience selection and drift independently of the other, so their evolutionary trajectories are not inevitably aligned and may directly conflict with one another [23-32]. Although the engulfed partner is capable of exerting selfish tendencies in some cases [33-35], there exist several mechanisms for the host to constrain the evolution of its symbionts [36-39]. In bacterial endosymbioses, this host-level constraint is often reflected in the genomic stasis of the bacterial partner. Endosymbiont genomes can remain stable in gene content and structure for tens [3], hundreds [12], or even thousands [5, 40] of millions of years, and we interpret this stability as a reflection of host-dominated evolution for the preservation of endosymbiont function.

However, secondary genome instability subsequent to this stasis is now recognized as relatively common, especially in mitochondria [41, 42]. Mitochondrial genomic instability manifests both as genome reduction [43, 44] that sometimes leads to outright genome loss [45-49], and genome fragmentation [50-53] that sometimes leads to massive genome expansion with little obvious functional change [54-57]. We suggest that what unites these starkly different outcomes is a shift away from the host-driven constraint of the endosymbiont genome towards (sometimes temporary) symbiont-driven instability. In cases of mitochondrial reduction and loss, the host ecology changes such that the function of the organelle is no longer needed and therefore no longer under selective constraint from the host [46-48]. For example, many eukaryotes that live in anaerobic environments no longer require the oxidative respiratory function of their mitochondria, so the genes for this process are free to be lost [44]. The forces promoting mitochondrial genome fragmentation and expansion are less clear, but these expansions sometimes seem to be associated with increases in mitochondrial mutation rates [55] and have been hypothesized to result from less efficacious host-level selection against slightly deleterious symbiont mutations [57, 58].

Depending on whether one takes a *Hodgkinia*- or cicada-centric perspective, the outcomes we report here could be interpreted either as a genome reductive or genome expansive process [7, 8]. From the perspective of individual *Hodgkinia* lineages, each circular molecule in *Magicicada* gets smaller after a split, eventually resulting in circles less than half the size and encoding fewer than ∼30% of the genes that were present on the already tiny ancestral *Hodgkinia* genome (Table S1)[6, 7]. This reductive process likely reflects the deletional mutation bias of bacteria [59], which in part explains the extremely small size of bacterial endosymbiont genomes in general [1]. In *Hodgkinia*, newly split lineages resemble a gene duplication event [7] that often results in one gene copy being pseudogenized and eventually lost entirely. *Hodgkinia*’s splitting and deletion process leads to individual circular molecules that resemble the extremely degraded genomes of mitochondria found in some eukaryotes. The idiosyncratic nature of these circles in closely related cicada species (Fig. 2C) is consistent with stochastic mutational loss and suggests a process with no particular goal or end point. But an important difference between cases of mitochondrial genome reduction and *Hodgkinia* from *Magicicada* is that the host ecology has not changed such that *Hodgkinia’*s functions are no longer required. The massive gene loss on individual *Hodgkinia* circles is likely only tolerable because, from the host’s perspective, the combined HGCs seem to have retained *Hodgkinia*’s overall nutritional contribution to the symbiosis (Fig. 2D). From the host perspective, this splitting and genome reductive process results in a combined *Hodgkinia “*genome” size over an order of magnitude larger than the ancestral single genome (Table 1).

In our view, the most interesting parallel to what we report here for *Hodgkinia* can be found in the mitochondrial genomes of the angiosperm genus *Silene* [55, 60]. Like many plants, some *Silene* mitochondrial genomes consist of a single “master circle” with multiple “subcircles” that arise primarily from recombination [61]. Other *Silene* species, though, have experienced dramatic increases in mitochondrial mutation rates, which seem to be accompanied by the expansion to dozens of enormous mitochondrial chromosomes [55]. These mitochondrial chromosomes, some encoding few or no detectable genes, can be rapidly lost or gained in closely related *Silene* lineages [60]. Like *Hodgkinia,* this diversity of genome expansion outcomes in closely related plant hosts is not accompanied by any detectable increase in functional capacity. We previously hypothesized that the increased complexity of *Hodgkinia* in *Magicicada* results from a similar increased effective mutation rate in *Hodgkinia* [8], with a conceptual modification related to lifecycle changes of the host cicada. While *Hodgkinia*’s inherent mutation rate may not be different in various cicada hosts, longer host lifecycles such as the 13-or 17-year lifecycle of *Magicicada* [62] may allow more symbiont generations and thus more *Hodgkinia* mutations per host lifecycle. We hypothesize that this increase in effective mutation rate enables *Hodgkinia*’s lineage splitting process and eventually results in stochastic differences between HGCs from different species groups at the level of genome structure (Fig. 2C). While *Hodgkinia* genes are (mostly) maintained in all HGCs, they are now present at wildly different abundances in different cicada species groups (Fig. 3). We hypothesize that lineage splitting and changes in gene dosages are either maladaptive or neutral for the host. The cicada does not benefit from *Hodgkinia* degeneration but must tolerate it because the cicada is wholly dependent on *Hodgkinia* for survival.

## Materials and Methods

### DNA extraction

Bacteriomes were dissected from a single male of *T. crassa*, a single female of *A. curvicosta* and *M. tredecim*, and two females of the remaining species. DNA was extracted from all dissected bacteriomes using a DNeasy Blood and Tissue kit (Qiagen catalog #69506). Extracted DNA was stored at -20C.

### Library preparation and sequencing

Genomic DNA from *M. tredecim* was sheared to an average fragment size of 550 base pairs using a Covaris E220. Sheared DNA was made into a sequencing library using the NEBNext Ultra DNA Library Prep Kit for Illumina (catalog #E7370S), according to the standard protocol. The library was sequenced at the University of Montana Genomics Core on a MiSeq benchtop sequencer with a v3 600 cycle kit.

Genomic DNA from *A. curvicosta* was sheared to an average size of 480 base pairs using a Covaris E220. Sheared DNA was made into a sequencing library using a TruSeq PCR-free kit (Illumina) and sequenced as ∼1/12 of a multiplexed lane at NGX Bio in San Francisco, CA using a HiSeq 2500 Rapid SBS kit (Illumina).

Genomic DNA from *T. crassa* was sheared to an average of 570 base pairs using a Covaris E220. Sheared DNA was made into a sequencing library using the NEBNext Ultra DNA Library Prep Kit for Illumina (catalog #E7370S), according to the standard protocol. The library was sequenced as ∼1/4 of a multiplexed lane at the University of Montana Genomics Core on a MiSeq benchtop sequencer with a v3 600 cycle kit.

Genomic DNA from *M. neotredecim, septendecim, tredecassini, cassini, tredecula*, and *septendecula* was sheared to an average of 500 base pairs using a Covaris E220. Sheared DNA was made into a sequencing library using the NEBNext Ultra DNA Library Prep Kit for Illumina (catalog #E7370S), according to the standard protocol. Libraries were sequenced on two lanes on a HiSeq 2500 with 250 cycles at the Johns Hopkins School of Medicine Genetic Resources Core Facility.

Genomic DNA from *M. neotredecim, septendecim, tredecassini, cassini, tredecula*, and *septendecula* was used for making libraries with a Nextera Mate Pair Sample Prep Kit (Illumina), according to the standard protocol. These libraries were sequenced on a single lane on a HiSeq 2500 with 100 cycles at the Case Western Reserve University Genomics Core Facility.

### Genome assembly and annotation

Raw reads were quality filtered using Trimmomatic version 0.35 [63]. Remaining reads were further filtered using fastq_quality_filter from FASTX version 0.0.13 (http://hannonlab.cshl.edu/fastx_toolkit/).

Assembly of the filtered reads was done using Spades version 3.6.2 [64], using kmer sizes 127, 151, 191, and 291, both individually as well as combined together. Putative *Hodgkinia* contigs were identified with TBLASTN 2.2.31+ [65] with an E-value cutoff of 10e-5 using previously annotated *Hodgkinia* genes as the query. To remove redundant contigs, all putatively *Hodgkinia* contigs were queried against themselves, and any contig >=97% identical to another over >= 80% of its length was considered redundant and removed. Any contigs with BLASTN E-values less than 10e-10 to the mitochondrial genome were also removed. Coverage of individual contigs was calculated by the total coverage at each base, divided by the length of the contig. Completely assembled *Hodgkinia* circles were identified based on sequence overlap on both ends of the contig. To identify putative circular contigs, filtered paired end and mate pair reads were mapped back to the assembly using BWA version 0.7.12-r1039 [66] with default parameters. Contigs were considered putatively circular if more than five read pairs mapped with one mate mapping in the first 10% of the contig, while its mate mapped in the last 10% of the contig. Putatively circular contigs were then closed when possible by PCR and Sanger sequencing.

Annotation of the *Hodgkinia* circles was done using a custom Python pipeline based around the Jackhmmer module of HMMER v. 3.1b2 [67], RNAmmer 1.2 [68], and Aragorn v1.2.36 [69]. Occasionally RNAmmer misannotated the 23S rRNA gene, so barrnap 0.6 [70] was used for corrections. The completely closed *Hodgkinia* circles were then checked manually for any long open reading frames that could contain missing genes.

### Phylogenetic analysis

Host phylogeny was reconstructed using RAxML v. 8.2.0 [71]based on manually inspected alignments of 15 mitochondrial genes (13 protein-coding and two rRNA) of the total length of 12744 bp, divided into four partitions corresponding to three codon positions and to rRNA genes. Rapid bootstraping (100 replicates) was used to estimate node support.

To construct individual gene phylogenies, homologous nucleotide sequences were translated into amino acids and aligned using mafft v. 7.221[74]. Visually inspected alignments were analyzed using RaxML v. 8.2.4 [71] using a PROTGAMMAWAG model of amino acid substitution and 100 bootstrap replicates. Trees were rooted using Aleeta-Tryella as outgroups (whenever they formed a single monophyletic clade), or alternatively on the longest branch separating well-supported clades that included species from all or most hosts in a comparison.

### Comparative *Hodgkinia* genome analysis

To compare the homology of HGC circles between cicada species, a *Hodgkinia* Similarity Index (HSI) score was calculated for all pairwise comparisons of all circles, as explained in Results. The pair with the highest HSI score was kept for each circle.

To determine relative coverage of all *Hodgkinia* genes, the coverage of all *Hodgkinia* genes was summed based on the coverage of the contig on which it was annotated. These abundance values were then normalized based on the most abundant gene. Principal coordinates analysis was done using the R package Vegan 2.4-3 [72].

## End Matter

### Author Contributions and Notes

Conceptualization, M.A.C. and J.P.M.; Methodology, M.A.C. and P.L.; Formal Analysis, M.A.C.; Investigation, M.A.C.; Resources, C.S. and J.P.M.; Data Curation, M.A.C. and C.S.; Writing – Original Draft, M.A.C.; Writing – Review & Editing, M.A.C., P.L., C.S., and J.P.M.; Visualization, M.A.C.; Supervision, J.P.M.; Funding Acquisition, J.P.M.

The authors declare no conflict of interest.

#### Acknowledgments

We thank all members of the McCutcheon lab for helpful discussion and comments, and Scott Miller for suggesting the use of the Jaccard Index. Funding for the sequencing and analysis was supported by National Science Foundation grants IOS-1256680 and IOS-1553529, and National Aeronautics and Space Administration Astrobiology Institute Award NNA15BB04A. Funding for cicada collecting was provided by NSF DEB-09-55849.

## Supplemental Information

**Figure S1:**
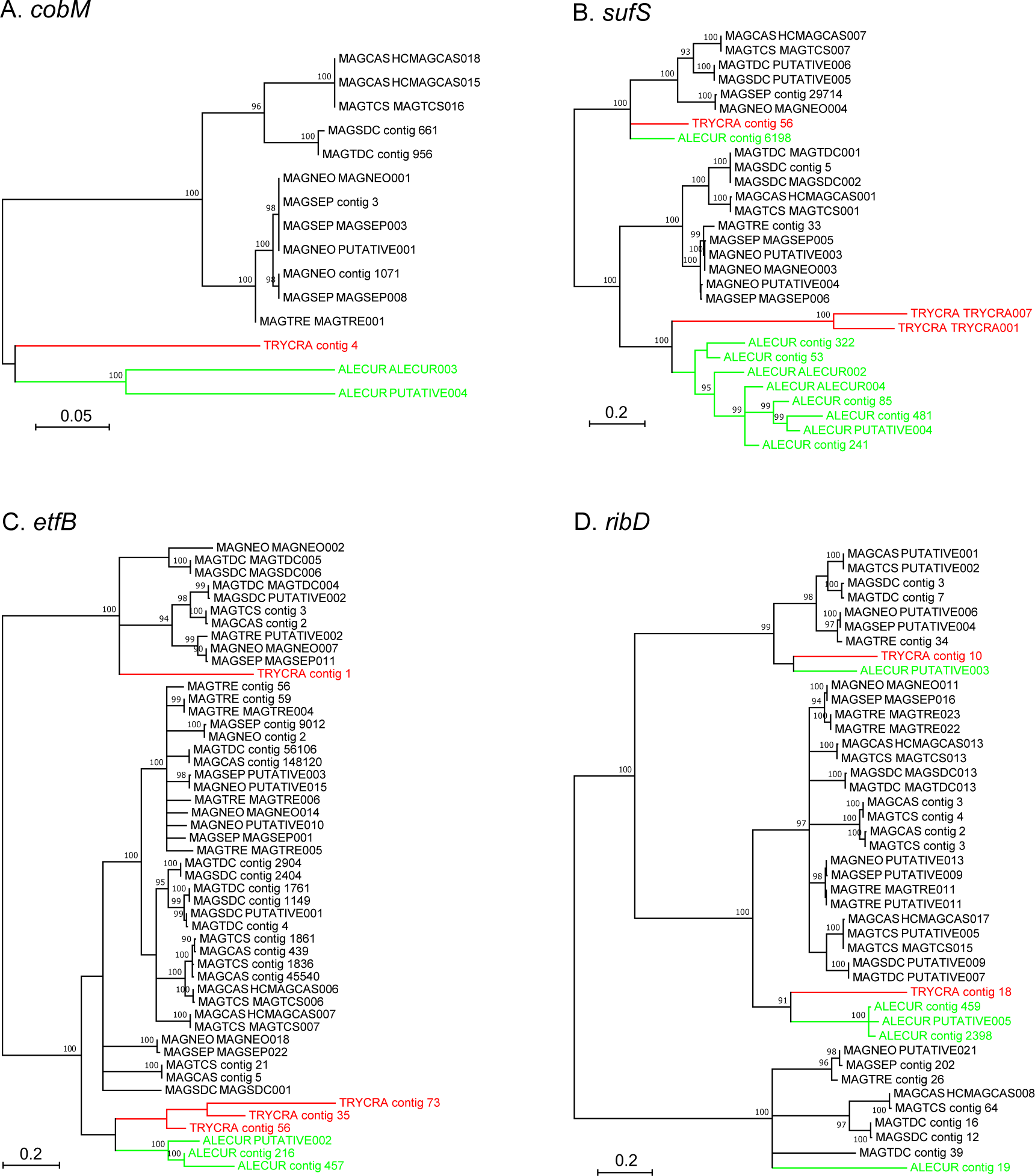
Phylogenetic trees showing ancient lineage splitting. Related to Figure 1. Maximum likelihood phylogenies of selected genes of *Hodgkinia* from *Magicicada* (black) and outgroups - *Aleeta curvicosta* (green) and *Tryella crassa* (red), based on amino acid sequences of all copies from these nine species. Bootstrap support values >90% are shown, nodes with support <70% were collapsed. Trees are rooted on longest branches. Tree A show patterns consistent with the presence of only a single gene copy in the last common ancestor, and independent splits in some of the derived host clades. Trees B-D show patterns consistent with the presence of at least two gene copies in the last common ancestor of *Magicicada* and at least one of the outgroups.

**Figure S2:**
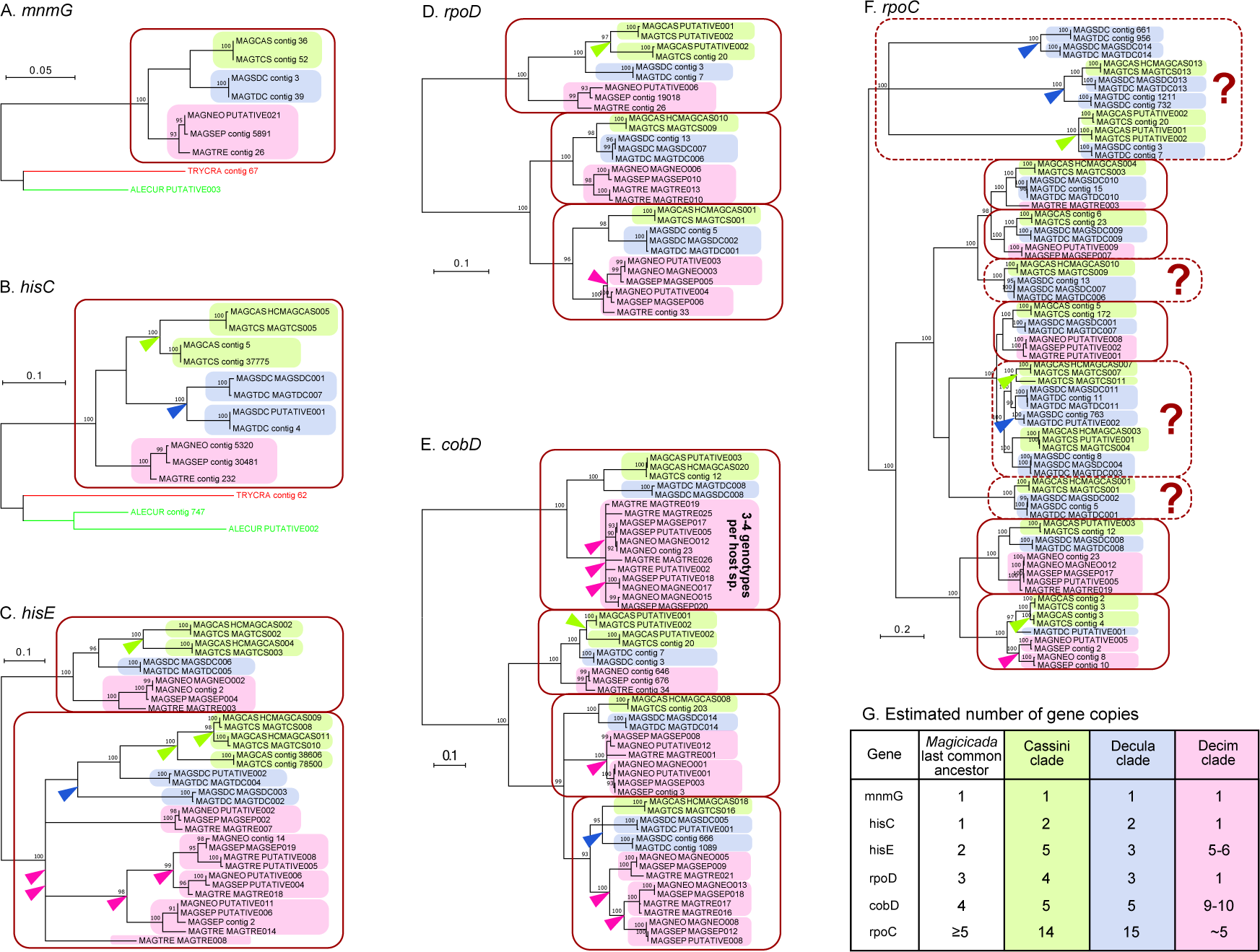
Phylogenetic trees showing numbers of distinct gene copies in the ancestor of *Magicicada*. Related to Figure 1. (A-F) Maximum likelihood phylogenies of selected genes of *Hodgkinia* from *Magicicada*, based on amino acid sequences of all copies from seven species. Bootstrap support values >90% are shown, nodes with support <70% were collapsed. Gene copies from Cassini, Decula and Decim species groups are indicated with green, blue and pink backgrounds, respectively. Solid maroon boxes indicate monophyletic clades that consist of gene copies from all species groups, and which are likely derived from a gene copy present in the last common ancestor. Dashed maroon boxes with question marks indicate monophyletic clades which may have derived from a gene copy or copies present in the common ancestor, but which do not include sequences from all species groups: it is possible that in the Decim species group, that gene copy was lost. Arrowheads indicate gene split events that took place after a species group diverged. Trees A-B are rooted using outgroups - *Aleeta* and *Tryella*; in other trees, the root has been placed on the longest branch between well-supported clades that included species from all or most hosts. (G) Numbers of distinct copies of selected genes in the last common ancestor of *Magicicada* and in extant species groups, estimated based on trees in panels A-F.

**Figure S3:**
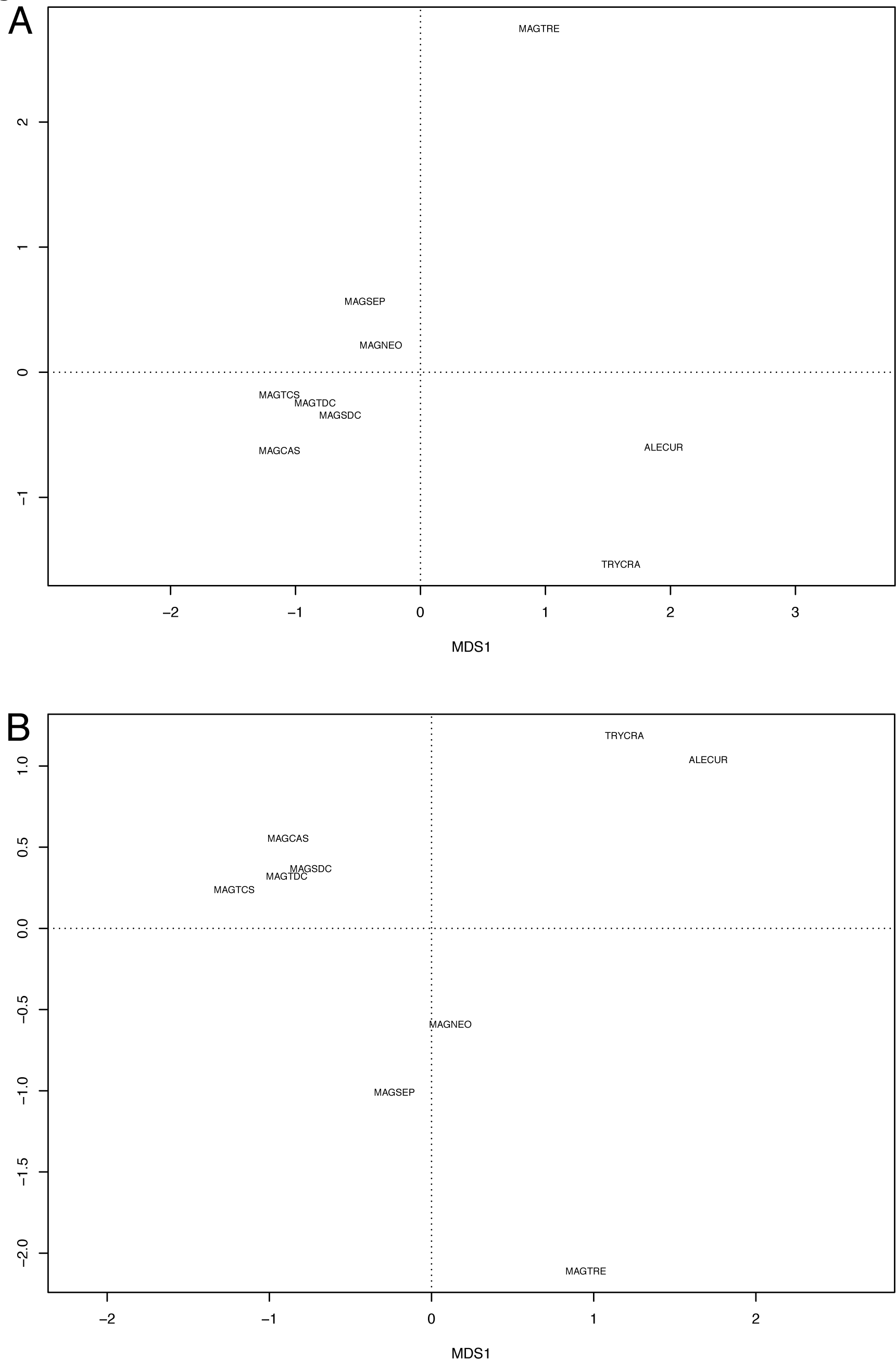
Principal coordinates chart of relative gene abundance. Related to
Figure 3. Charts from the principal coordinates analysis of relative gene abundance from Figure 3. A) All genes present in any species. B) Only genes present in all species. Species abbreviations are taken from Table 1.

| Species | Scaffold | Scaffold length | Number of genes | Coding density (%) | GC content (%) | Average coverage |
| --- | --- | --- | --- | --- | --- | --- |
| *M. cassini* | MAGCAS001 | 69473 | 17 | 31 | 34 | 2535 |
| *M. cassini* | MAGCAS002 | 61653 | 13 | 20.7 | 29.9 | 1625 |
| *M. cassini* | MAGCAS003 | 52116 | 6 | 21.5 | 27.1 | 1263 |
| *M. cassini* | MAGCAS004 | 48995 | 18 | 43.6 | 31.9 | 2219 |
| *M. cassini* | MAGCAS005 | 45702 | 9 | 25.2 | 31.4 | 515 |
| *M. cassini* | MAGCAS006 | 41268 | 14 | 39.2 | 32.1 | 580 |
| *M. cassini* | MAGCAS007 | 35613 | 8 | 48.5 | 29.9 | 216 |
| *M. cassini* | MAGCAS008 | 33827 | 5 | 21.9 | 38.6 | 682 |
| *M. cassini* | MAGCAS009 | 32518 | 4 | 17.6 | 35.3 | 158 |
| *M. cassini* | MAGCAS010 | 31635 | 4 | 31.7 | 33.6 | 509 |
| *M. cassini* | MAGCAS011 | 29608 | 4 | 18 | 35.2 | 673 |
| *M. cassini* | MAGCAS012 | 18833 | 4 | 29.9 | 38.1 | 502 |
| *M. cassini* | MAGCAS013 | 18274 | 4 | 52.1 | 36.4 | 944 |
| *M. cassini* | MAGCAS014 | 18203 | 6 | 45.2 | 36.4 | 2231 |
| *M. cassini* | MAGCAS015 | 17469 | 3 | 12.9 | 36.5 | 1141 |
| *M. cassini* | MAGCAS016 | 16309 | 4 | 15.3 | 28.3 | 182 |
| *M. cassini* | MAGCAS017 | 15578 | 3 | 18.9 | 36.1 | 666 |
| *M. cassini* | MAGCAS018 | 13935 | 4 | 40.8 | 34.6 | 1032 |
| *M. cassini* | MAGCAS019 | 9976 | 4 | 59.9 | 45.2 | 689 |
| *M. cassini* | MAGCAS020 | 8941 | 1 | 10.7 | 23.1 | 553 |
| *M. cassini* | MAGCAS021 | 8091 | 4 | 62.2 | 45.3 | 1119 |
| *M. cassini* | MAGCAS022 | 5678 | 3 | 41.4 | 34 | 544 |
| *M. cassini* | MAGCAS023 | 1057 | 1 | 60.1 | 29.7 | 2642 |
| *M. cassini* | MAGCAS024 | 748 | 1 | 62.4 | 32.5 | 101 |
| *M. cassini* | PUTATIVE001 | 46023 | 11 | 34.3 | 29.7 | 870 |
| *M. cassini* | PUTATIVE002 | 28503 | 6 | 39.4 | 31.3 | 305 |
| *M. cassini* | PUTATIVE003 | 27789 | 6 | 37.5 | 25.4 | 836 |
| *M. cassini* | PUTATIVE004 | 21863 | 2 | 9.3 | 31 | 2590 |
| *M. cassini* | PUTATIVE006 | 14618 | 1 | 10.2 | 31.5 | 204 |
| *M. tredecassini* | MAGTCS001 | 69472 | 17 | 30.7 | 34 | 1450 |
| *M. tredecassini* | MAGTCS002 | 61641 | 13 | 20.8 | 29.9 | 1646 |
| *M. tredecassini* | MAGTCS003 | 56215 | 21 | 47.9 | 33.1 | 2524 |
| *M. tredecassini* | MAGTCS004 | 52125 | 6 | 21.6 | 27 | 753 |
| *M. tredecassini* | MAGTCS005 | 45699 | 10 | 28 | 31.5 | 340 |
| *M. tredecassini* | MAGTCS006 | 41278 | 14 | 39.1 | 32.2 | 80 |
| *M. tredecassini* | MAGTCS007 | 35614 | 8 | 48.5 | 29.9 | 129 |
| *M. tredecassini* | MAGTCS008 | 32520 | 3 | 15.6 | 35.4 | 502 |
| *M. tredecassini* | MAGTCS009 | 31636 | 4 | 31.7 | 33.6 | 237 |
| *M. tredecassini* | MAGTCS010 | 29317 | 4 | 18.2 | 35 | 339 |
| *M. tredecassini* | MAGTCS011 | 22416 | 17 | 73.3 | 31.1 | 4356 |
| *M. tredecassini* | MAGTCS012 | 18834 | 4 | 29.9 | 38.2 | 190 |
| *M. tredecassini* | MAGTCS013 | 18271 | 4 | 52.1 | 36.4 | 500 |
| *M. tredecassini* | MAGTCS014 | 16306 | 3 | 10.7 | 28.4 | 183 |
| *M. tredecassini* | MAGTCS015 | 15579 | 3 | 18.9 | 36 | 549 |
| *M. tredecassini* | MAGTCS016 | 13934 | 4 | 40.8 | 34.5 | 996 |
| *M. tredecassini* | MAGTCS017 | 5680 | 3 | 40.7 | 33.9 | 172 |
| *M. tredecassini* | MAGTCS018 | 1057 | 1 | 68.6 | 29.8 | 1424 |
| *M. tredecassini* | MAGTCS019 | 748 | 1 | 63.2 | 32.5 | 83 |
| *M. tredecassini* | MAGTCS020 | 694 | 1 | 94.1 | 25.4 | 54 |
| *M. tredecassini* | PUTATIVE001 | 58824 | 6 | 19.1 | 27.1 | 738 |
| *M. tredecassini* | PUTATIVE002 | 46026 | 10 | 33 | 29.7 | 806 |
| *M. tredecassini* | PUTATIVE003 | 21389 | 2 | 9.5 | 30.9 | 1839 |
| *M. tredecassini* | PUTATIVE005 | 15824 | 3 | 18.6 | 35.9 | 552 |
| *M. tredecassini* | PUTATIVE006 | 15349 | 3 | 17.9 | 27.7 | 2623 |
| *M. tredecassini* | PUTATIVE007 | 11443 | 3 | 45 | 38.5 | 1237 |
| *M. septendecula* | MAGSDC001 | 70504 | 27 | 36.7 | 28.6 | 2344 |
| *M. septendecula* | MAGSDC002 | 70413 | 16 | 28.5 | 31.9 | 290 |
| *M. septendecula* | MAGSDC003 | 53932 | 6 | 14.8 | 25.9 | 664 |
| *M. septendecula* | MAGSDC004 | 51941 | 6 | 21.4 | 24 | 311 |
| *M. septendecula* | MAGSDC005 | 41633 | 5 | 11.5 | 28.6 | 1476 |
| *M. septendecula* | MAGSDC006 | 34671 | 17 | 43.4 | 32.8 | 2072 |
| *M. septendecula* | MAGSDC007 | 31673 | 4 | 31.7 | 32.4 | 196 |
| *M. septendecula* | MAGSDC008 | 29204 | 7 | 29.4 | 27.7 | 1293 |
| *M. septendecula* | MAGSDC009 | 29090 | 7 | 51 | 34 | 448 |
| *M. septendecula* | MAGSDC010 | 29000 | 7 | 50.8 | 31.9 | 1642 |
| *M. septendecula* | MAGSDC011 | 22533 | 9 | 48.5 | 30.1 | 1820 |
| *M. septendecula* | MAGSDC012 | 18788 | 7 | 46 | 37.6 | 495 |
| *M. septendecula* | MAGSDC013 | 17780 | 4 | 51.2 | 34.4 | 174 |
| *M. septendecula* | MAGSDC014 | 13841 | 3 | 60.8 | 30.6 | 388 |
| *M. septendecula* | MAGSDC015 | 10837 | 4 | 53.8 | 46.7 | 1520 |
| *M. septendecula* | MAGSDC016 | 8802 | 2 | 47.8 | 43 | 493 |
| *M. septendecula* | MAGSDC017 | 7860 | 2 | 53.7 | 47.3 | 331 |
| *M. septendecula* | MAGSDC018 | 1107 | 0 | 0 | 27.4 | 1637 |
| *M. septendecula* | MAGSDC019 | 748 | 1 | 63.2 | 29.4 | 72 |
| *M. septendecula* | MAGSDC020 | 692 | 1 | 93.9 | 26.3 | 17 |
| *M. septendecula* | PUTATIVE001 | 66259 | 20 | 26.9 | 28.7 | 456 |
| *M. septendecula* | PUTATIVE002 | 31705 | 6 | 12.8 | 31.2 | 1220 |
| *M. septendecula* | PUTATIVE003 | 20842 | 4 | 10.3 | 28.2 | 532 |
| *M. septendecula* | PUTATIVE005 | 14087 | 8 | 63.8 | 24.7 | 1702 |
| *M. septendecula* | PUTATIVE006 | 11820 | 9 | 36.6 | 29.1 | 4455 |
| *M. septendecula* | PUTATIVE008 | 10372 | 3 | 11.3 | 31.4 | 914 |
| *M. septendecula* | PUTATIVE009 | 10281 | 3 | 28.6 | 37.5 | 1636 |
| *M. tredecula* | MAGTDC001 | 70424 | 15 | 29.3 | 31.9 | 465 |
| *M. tredecula* | MAGTDC002 | 54074 | 6 | 20 | 25.9 | 182 |
| *M. tredecula* | MAGTDC003 | 51873 | 6 | 21.4 | 24 | 154 |
| *M. tredecula* | MAGTDC004 | 47884 | 11 | 20.1 | 32.7 | 735 |
| *M. tredecula* | MAGTDC005 | 34671 | 17 | 43.4 | 32.8 | 1702 |
| *M. tredecula* | MAGTDC006 | 31664 | 4 | 31.7 | 32.4 | 479 |
| *M. tredecula* | MAGTDC007 | 29603 | 12 | 44.9 | 28.7 | 2787 |
| *M. tredecula* | MAGTDC008 | 29201 | 7 | 29.3 | 27.8 | 500 |
| *M. tredecula* | MAGTDC009 | 29085 | 7 | 51.1 | 34 | 480 |
| *M. tredecula* | MAGTDC010 | 29002 | 7 | 50.8 | 31.9 | 574 |
| *M. tredecula* | MAGTDC011 | 22533 | 10 | 57.5 | 30.1 | 571 |
| *M. tredecula* | MAGTDC012 | 18787 | 7 | 46.1 | 37.6 | 570 |
| *M. tredecula* | MAGTDC013 | 17780 | 4 | 51.2 | 34.4 | 131 |
| *M. tredecula* | MAGTDC014 | 13840 | 3 | 61.2 | 30.6 | 251 |
| *M. tredecula* | MAGTDC015 | 10839 | 4 | 53.8 | 46.8 | 1096 |
| *M. tredecula* | MAGTDC016 | 8803 | 2 | 47.8 | 43 | 640 |
| *M. tredecula* | MAGTDC017 | 7866 | 2 | 53.6 | 47.3 | 633 |
| *M. tredecula* | MAGTDC018 | 1108 | 1 | 65.4 | 27.8 | 514 |
| *M. tredecula* | MAGTDC019 | 748 | 1 | 58.8 | 29.3 | 27 |
| *M. tredecula* | MAGTDC020 | 692 | 1 | 93.9 | 26.3 | 16 |
| *M. tredecula* | PUTATIVE001 | 63118 | 12 | 24.1 | 30.4 | 856 |
| *M. tredecula* | PUTATIVE002 | 35545 | 17 | 63.7 | 30.7 | 2467 |
| *M. tredecula* | PUTATIVE003 | 20858 | 4 | 10.3 | 28.1 | 335 |
| *M. tredecula* | PUTATIVE004 | 16213 | 2 | 10.2 | 30.3 | 780 |
| *M. tredecula* | PUTATIVE005 | 13731 | 3 | 33.7 | 39.1 | 459 |
| *M. tredecula* | PUTATIVE006 | 13664 | 8 | 65.8 | 24.5 | 734 |
| *M. tredecula* | PUTATIVE007 | 10286 | 3 | 28.6 | 37.5 | 809 |
| *M. neotredecim* | MAGNEO001 | 61256 | 7 | 9 | 24.9 | 134 |
| *M. neotredecim* | MAGNEO002 | 57172 | 21 | 38.7 | 29.9 | 257 |
| *M. neotredecim* | MAGNEO003 | 56251 | 12 | 21.2 | 29.6 | 180 |
| *M. neotredecim* | MAGNEO004 | 35088 | 11 | 27.4 | 23.7 | 249 |
| *M. neotredecim* | MAGNEO005 | 34472 | 4 | 16.5 | 31 | 366 |
| *M. neotredecim* | MAGNEO006 | 34053 | 3 | 7.4 | 29.9 | 187 |
| *M. neotredecim* | MAGNEO007 | 27670 | 4 | 9.1 | 28.1 | 120 |
| *M. neotredecim* | MAGNEO008 | 23724 | 5 | 23.2 | 32.3 | 155 |
| *M. neotredecim* | MAGNEO009 | 22209 | 4 | 32.2 | 35.8 | 161 |
| *M. neotredeceim* | MAGNEO010 | 21502 | 2 | 9 | 28.4 | 103 |
| *M. neotredeceim* | MAGNEO011 | 19133 | 2 | 8 | 32.5 | 148 |
| *M. neotredeceim* | MAGNEO012 | 18932 | 4 | 48.6 | 24.4 | 147 |
| *M. neotredeceim* | MAGNEO013 | 17725 | 1 | 5.3 | 27.2 | 173 |
| *M. neotredeceim* | MAGNEO014 | 12466 | 4 | 21.4 | 26.6 | 164 |
| *M. neotredeceim* | MAGNEO015 | 11399 | 2 | 12.5 | 21.9 | 81 |
| *M. neotredeceim* | MAGNEO016 | 11218 | 1 | 3.8 | 24.7 | 64 |
| *M. neotredeceim* | MAGNEO017 | 10677 | 2 | 13.4 | 22 | 530 |
| *M. neotredeceim* | MAGNEO018 | 3222 | 2 | 57.4 | 25.2 | 152 |
| *M. neotredeceim* | MAGNEO019 | 1100 | 1 | 66.2 | 26.7 | 715 |
| *M. neotredeceim* | MAGNEO020 | 748 | 1 | 60 | 30.9 | 81 |
| *M. neotredeceim* | PUTATIVE001 | 64450 | 7 | 8.6 | 24.9 | 126 |
| *M. neotredeceim* | PUTATIVE002 | 59782 | 6 | 12.6 | 23.3 | 53 |
| *M. neotredeceim* | PUTATIVE003 | 55986 | 12 | 20.9 | 29.5 | 183 |
| *M. neotredeceim* | PUTATIVE004 | 54126 | 8 | 12.8 | 29.1 | 155 |
| *M. neotredeceim* | PUTATIVE005 | 53050 | 9 | 25.9 | 29.6 | 347 |
| *M. neotredeceim* | PUTATIVE006 | 41569 | 7 | 12.8 | 27.5 | 184 |
| *M. neotredeceim* | PUTATIVE007 | 30380 | 1 | 4.3 | 24 | 55 |
| *M. neotredeceim* | PUTATIVE008 | 30105 | 13 | 41.7 | 25.2 | 363 |
| *M. neotredeceim* | PUTATIVE009 | 28933 | 6 | 38.1 | 31.2 | 306 |
| *M. neotredeceim* | PUTATIVE010 | 27031 | 10 | 30.3 | 25.9 | 402 |
| *M. neotredeceim* | PUTATIVE011 | 26196 | 3 | 8 | 27.6 | 270 |
| *M. neotredeceim* | PUTATIVE012 | 21868 | 1 | 4.2 | 26.4 | 5 |
| *M. neotredeceim* | PUTATIVE013 | 20101 | 4 | 21.2 | 34 | 256 |
| *M. neotredeceim* | PUTATIVE014 | 18183 | 5 | 29.6 | 27.6 | 257 |
| *M. neotredeceim* | PUTATIVE015 | 17100 | 6 | 32.7 | 27.5 | 748 |
| *M. neotredeceim* | PUTATIVE016 | 15929 | 1 | 5.7 | 27.9 | 289 |
| *M. neotredeceim* | PUTATIVE018 | 14006 | 3 | 30 | 29.5 | 211 |
| *M. neotredeceim* | PUTATIVE019 | 12258 | 3 | 43.6 | 32.8 | 449 |
| *M. neotredeceim* | PUTATIVE020 | 10927 | 4 | 39.4 | 30.8 | 688 |
| *M. neotredeceim* | PUTATIVE021 | 10855 | 4 | 41.6 | 32.4 | 270 |
| *M. neotredeceim* | PUTATIVE022 | 10248 | 2 | 8.4 | 25.9 | 166 |
| *M. septendecim* | MAGSEP001 | 63626 | 22 | 27 | 27.2 | 724 |
| *M. septendecim* | MAGSEP002 | 61848 | 7 | 13.7 | 23.5 | 425 |
| *M. septendecim* | MAGSEP003 | 61285 | 7 | 8.8 | 24.9 | 534 |
| *M. septendecim* | MAGSEP004 | 57162 | 20 | 37.6 | 29.9 | 2101 |
| *M. septendecim* | MAGSEP005 | 56298 | 12 | 21.2 | 29.5 | 1285 |
| *M. septendecim* | MAGSEP006 | 56278 | 9 | 17.5 | 29.4 | 176 |
| *M. septendecim* | MAGSEP007 | 48066 | 12 | 37.1 | 30.2 | 863 |
| *M. septendecim* | MAGSEP008 | 43440 | 3 | 7.2 | 25.4 | 73 |
| *M. septendecim* | MAGSEP009 | 34489 | 4 | 16.5 | 30.9 | 781 |
| *M. septendecim* | MAGSEP010 | 33034 | 3 | 7.7 | 30.1 | 425 |
| *M. septendecim* | MAGSEP011 | 28244 | 4 | 8.9 | 28.3 | 487 |
| *M. septendecim* | MAGSEP012 | 22960 | 5 | 24.3 | 32 | 208 |
| *M. septendecim* | MAGSEP013 | 22204 | 4 | 25 | 35.8 | 348 |
| *M. septendecim* | MAGSEP014 | 21903 | 3 | 12.5 | 28.6 | 835 |
| *M. septendecim* | MAGSEP015 | 21496 | 2 | 9 | 28.4 | 440 |
| *M. septendecim* | MAGSEP016 | 19133 | 2 | 8 | 32.4 | 351 |
| *M. septendecim* | MAGSEP017 | 18932 | 4 | 48.6 | 24.3 | 350 |
| *M. septendecim* | MAGSEP018 | 17726 | 1 | 5.3 | 27.2 | 231 |
| *M. septendecim* | MAGSEP019 | 15979 | 3 | 30.4 | 35.8 | 697 |
| *M. septendecim* | MAGSEP020 | 11397 | 2 | 12.5 | 22 | 223 |
| *M. septendecim* | MAGSEP021 | 11166 | 2 | 38.1 | 42.4 | 385 |
| *M. septendecim* | MAGSEP022 | 3224 | 2 | 55.3 | 25.1 | 54 |
| *M. septendecim* | MAGSEP023 | 1099 | 1 | 66.2 | 26.8 | 753 |
| *M. septendecim* | PUTATIVE001 | 49706 | 4 | 11.8 | 25.7 | 360 |
| *M. septendecim* | PUTATIVE002 | 41910 | 19 | 46.4 | 26.2 | 1773 |
| *M. septendecim* | PUTATIVE003 | 33062 | 11 | 30.7 | 27.5 | 1566 |
| *M. septendecim* | PUTATIVE004 | 30610 | 5 | 10.2 | 26.1 | 780 |
| *M. septendecim* | PUTATIVE005 | 30280 | 6 | 35.3 | 23.4 | 289 |
| *M. septendecim* | PUTATIVE006 | 26244 | 3 | 7.9 | 27.5 | 978 |
| *M. septendecim* | PUTATIVE007 | 23391 | 5 | 18.7 | 28 | 2022 |
| *M. septendecim* | PUTATIVE008 | 23205 | 5 | 23.9 | 32 | 215 |
| *M. septendecim* | PUTATIVE009 | 20105 | 4 | 21.2 | 34 | 856 |
| *M. septendecim* | PUTATIVE010 | 18957 | 4 | 9.9 | 22.9 | 684 |
| *M. septendecim* | PUTATIVE011 | 18955 | 3 | 7.3 | 22.9 | 216 |
| *M. septendecim* | PUTATIVE013 | 15938 | 3 | 6.3 | 24.9 | 1209 |
| *M. septendecim* | PUTATIVE014 | 14676 | 3 | 10.7 | 24.5 | 351 |
| *M. septendecim* | PUTATIVE015 | 13624 | 6 | 20.6 | 26.7 | 836 |
| *M. septendecim* | PUTATIVE017 | 10118 | 3 | 36.1 | 32.2 | 736 |
| *M. septendecim* | PUTATIVE018 | 10107 | 2 | 14.1 | 21.8 | 857 |
| *M. tredecim* | MAGTRE001 | 61247 | 7 | 9 | 24.7 | 384 |
| *M. tredecim* | MAGTRE002 | 59026 | 9 | 16.2 | 30.4 | 504 |
| *M. tredecim* | MAGTRE003 | 58092 | 21 | 45.1 | 29.8 | 1392 |
| *M. tredecim* | MAGTRE004 | 55954 | 19 | 29.2 | 27.8 | 1419 |
| *M. tredecim* | MAGTRE005 | 52863 | 12 | 22.2 | 26.3 | 925 |
| *M. tredecim* | MAGTRE006 | 46330 | 19 | 32 | 27.5 | 1043 |
| *M. tredecim* | MAGTRE007 | 43811 | 5 | 8.3 | 22.9 | 233 |
| *M. tredecim* | MAGTRE008 | 41390 | 10 | 21.4 | 32.3 | 503 |
| *M. tredecim* | MAGTRE009 | 40594 | 2 | 5 | 28.3 | 116 |
| *M. tredecim* | MAGTRE010 | 33207 | 3 | 7.6 | 30 | 155 |
| *M. tredecim* | MAGTRE011 | 32598 | 4 | 19.1 | 32.6 | 291 |
| *M. tredecim* | MAGTRE012 | 27904 | 6 | 21.7 | 35.7 | 401 |
| *M. tredecim* | MAGTRE013 | 25773 | 3 | 9.7 | 29.4 | 184 |
| *M. tredecim* | MAGTRE014 | 25212 | 4 | 14.3 | 27.4 | 421 |
| *M. tredecim* | MAGTRE015 | 24039 | 9 | 44.5 | 30.6 | 425 |
| *M. tredecim* | MAGTRE016 | 23178 | 1 | 4 | 25.6 | 57 |
| *M. tredecim* | MAGTRE017 | 23156 | 1 | 4 | 25.7 | 19 |
| *M. tredecim* | MAGTRE018 | 19535 | 3 | 8.6 | 31.7 | 262 |
| *M. tredecim* | MAGTRE019 | 19044 | 3 | 45.1 | 24.2 | 144 |
| *M. tredecim* | MAGTRE020 | 18064 | 2 | 25.5 | 24 | 256 |
| *M. tredecim* | MAGTRE021 | 13768 | 2 | 12.9 | 27.1 | 184 |
| *M. tredecim* | MAGTRE022 | 12894 | 1 | 7.2 | 32.2 | 157 |
| *M. tredecim* | MAGTRE023 | 12893 | 2 | 11.8 | 32.4 | 115 |
| *M. tredecim* | MAGTRE024 | 12600 | 3 | 16.1 | 25.7 | 1136 |
| *M. tredecim* | MAGTRE025 | 11445 | 2 | 12.4 | 21.6 | 175 |
| *M. tredecim* | MAGTRE026 | 11402 | 1 | 7.6 | 21.5 | 229 |
| *M. tredecim* | MAGTRE027 | 11191 | 2 | 16 | 33.7 | 223 |
| *M. tredecim* | MAGTRE028 | 11158 | 2 | 38.1 | 42.6 | 344 |
| *M. tredecim* | MAGTRE029 | 10693 | 5 | 54.2 | 40.2 | 99 |
| *M. tredecim* | MAGTRE030 | 9940 | 1 | 7.4 | 30.3 | 84 |
| *M. tredecim* | PUTATIVE001 | 43403 | 20 | 46 | 26 | 1480 |
| *M. tredecim* | PUTATIVE002 | 43134 | 5 | 7.5 | 28.5 | 184 |
| *M. tredecim* | PUTATIVE003 | 39854 | 3 | 4.9 | 31.3 | 150 |
| *M. tredecim* | PUTATIVE004 | 31153 | 3 | 11.8 | 28.3 | 220 |
| *M. tredecim* | PUTATIVE005 | 22075 | 4 | 22.2 | 24.4 | 91 |
| *M. tredecim* | PUTATIVE006 | 19049 | 2 | 21.7 | 27.9 | 236 |
| *M. tredecim* | PUTATIVE008 | 15164 | 4 | 33.6 | 35.7 | 84 |
| *M. tredecim* | PUTATIVE009 | 13153 | 0 | 0 | 26.5 | 192 |
| *M. tredecim* | PUTATIVE010 | 11299 | 8 | 54.4 | 24.5 | 4373 |
| *M. tredecim* | PUTATIVE011 | 11004 | 3 | 16.5 | 34.5 | 237 |
| *M. tredecim* | PUTATIVE012 | 10250 | 3 | 27.6 | 39.5 | 151 |
| *M. tredecim* | PUTATIVE013 | 8821 | 2 | 9.5 | 25.4 | 1121 |
| *T. crassa* | TRYCRA001 | 64739 | 8 | 8.3 | 20.9 | 140 |
| *T. crassa* | TRYCRA002 | 38749 | 4 | 11.4 | 21.1 | 68 |
| *T. crassa* | TRYCRA003 | 25142 | 3 | 7.6 | 24.2 | 76 |
| *T. crassa* | TRYCRA004 | 23146 | 4 | 7.1 | 20.4 | 51 |
| *T. crassa* | TRYCRA005 | 23137 | 1 | 1.8 | 20.3 | 121 |
| *T. crassa* | TRYCRA006 | 14624 | 2 | 10.1 | 20.6 | 58 |
| *T. crassa* | TRYCRA007 | 14492 | 2 | 11.8 | 15.9 | 89 |
| *T. crassa* | TRYCRA008 | 12260 | 1 | 2.9 | 28.8 | 38 |
| *T. crassa* | TRYCRA009 | 11775 | 1 | 10.9 | 19.8 | 14 |
| *T. crassa* | TRYCRA010 | 11711 | 3 | 38.9 | 44.1 | 264 |
| *T. crassa* | TRYCRA011 | 9758 | 1 | 9.6 | 14.4 | 39 |
| *T. crassa* | TRYCRA012 | 9446 | 1 | 9 | 13.8 | 43 |
| *T. crassa* | TRYCRA013 | 8867 | 2 | 10.6 | 14.4 | 93 |
| *T. crassa* | TRYCRA014 | 2151 | 2 | 88.3 | 22.5 | 20 |
| *A. curvica* | ALECUR001 | 46134 | 2 | 2.6 | 24 | 15 |
| *A. curvica* | ALECUR002 | 32844 | 3 | 12.2 | 32.6 | 16 |
| *A. curvica* | ALECUR003 | 23214 | 5 | 17.1 | 28.1 | 41 |
| *A. curvica* | ALECUR004 | 20530 | 3 | 19.6 | 33.1 | 17 |
| *A. curvica* | ALECUR005 | 7857 | 1 | 24.3 | 35.9 | 7 |
| *A. curvica* | PUTATIVE001 | 59192 | 5 | 6.1 | 28.1 | 41 |
| *A. curvica* | PUTATIVE002 | 55340 | 14 | 21.7 | 24.9 | 246 |
| *A. curvica* | PUTATIVE003 | 39946 | 9 | 18.4 | 30.1 | 18 |
| *A. curvica* | PUTATIVE004 | 34099 | 6 | 19.9 | 28.3 | 36 |
| *A. curvica* | PUTATIVE005 | 17753 | 2 | 7.1 | 35.5 | 43 |
| *A. curvica* | PUTATIVE006 | 16143 | 10 | 75.1 | 32.7 | 648 |

